# Neuronal oversight of germline small RNAs prevents heat-induced sterility in lab-domesticated *Caenorhabditis elegans*

**DOI:** 10.1101/2025.08.07.669182

**Authors:** Chee Kiang Ewe, Hanna Achache, Shir Weiss, Anna Mogilevskaya, Myriam Valenski, Guy Teichman, Sarit Anava, Hila Gingold, Rachel Posner, Olga Antonova, Yonatan B. Tzur, Oded Rechavi

## Abstract

Thermal pollution, whether local or driven by global warming, threatens biodiversity in part through its detrimental effects on reproduction. Non-coding small RNAs (sRNAs) are crucial for maintaining germline developmental robustness under heat stress. Remarkably, we uncovered that neuronal sRNAs regulate germ cells’ thermotolerance, affecting both spermatogenic and oogenic germline in a cell non-autonomous manner. Furthermore, we demonstrate that an oxygen-sensing neural circuit antagonizes germline maintenance, likely reflecting the nematode’s innate association of reduced oxygen levels with food availability and reproductive permissive environments. Finally, we provide evidence that sRNAs buffer the negative consequences of laboratory domestication, which otherwise cause the laboratory strain to lose germline integrity and become sterile at elevated temperatures. Hence, our findings reveal that mere sensory perception, independent of direct environmental change, modulates germline integrity through sRNA pathways, highlighting a novel mechanism by which neural circuits integrate environmental information to safeguard reproductive fitness in fluctuating environments.

## Main Text

*“We are going to die, and that makes us the lucky ones. Most people are never going to die because they are never going to be born.”* Underlying Richard Dawkins’ beautiful words is an important reminder: intricate molecular machinery has evolved to ensure germline immortality – but this is not to be taken for granted. Reproductive health is strongly influenced by environmental factors. Mass extinctions are happening at unprecedented scale, driven in large part by rising global temperatures that cause widespread fertility loss. Climate change is linked to declining male and female fertility from plants to mammals, posing a significant challenge for future generations (*1*, *2*). How do organisms endure and adapt in such difficult times?

Non-coding small RNAs (sRNAs), including siRNAs, miRNAs, and piRNAs, together with Argonautes (AGOs), play conserved roles in germline development and fertility (*3*). In flies and mammals, loss of PIWI proteins leads to germ cell loss and transposon de-repression (*4–6*). In mice, endo-siRNAs are critical for oogenesis, while miRNAs and piRNA are essential for multiple aspects of spermatogenesis (*7–11*). Similarly, in the nematode *C. elegans*, sRNAs play key roles in germ cell development (*12–17*). Given the potential of sRNA/AGO pathways to drive plastic gene regulatory programs, it is of great interest to understand how they regulate fecundity in changing environments.

Here, we show that neuron-to-germline communication plays an important role in regulating germline development under heat stress in laboratory-domesticated *C. elegans*. Specifically, we found that the oxygen-sensing neural circuit – a recurring target of environmental adaptation – modulates sRNA-mediated germline development in the presence of heat stress. Our results further reveal the molecular basis by which environmental sensory perception – despite no change in absolute oxygen levels – influences developmental robustness of germ cells through neuropeptide signaling and highlight how epigenetic mechanisms may buffer the negative effects of laboratory-derived mutations on reproductive fitness, particularly in stressful conditions.

## Endo-siRNAs protect sperm thermotolerance

TRBP2 is a dsRNA-binding protein essential for Dicer-mediated miRNA production in vertebrates (*18*). In flies, its homolog R2D2 and Loqs-PD, together with Dicer, facilitate siRNA loading onto RISC (*19*, *20*). In *C. elegans*, the TRBP2 homolog RDE-4 is critical for exogenous RNAi and antiviral defense by binding long dsRNAs and interacting with DCR-1/Dicer, DRH-1/RIC-I, and the AGO RDE-1 (*21–24*). RDE-4 is also required for the biogenesis of endogenous siRNAs and ensures their proper loading onto AGOs (*25*, *26*). Central to this study, previous work uncovered an important role of RDE-4 in embryogenesis during heat stress, with the loss-of-function mutants exhibiting temperature-sensitive developmental defects and laying arrested embryos at 25 °C (*27*).

Here, we similarly found that *rde-4*(-) mutants are fertile at 15 °C and 20 °C; however, our results show that, at 25 °C, they lay unfertilized oocytes – large, dark, rounded cells with prominent nuclei – that are readily distinguishable from unhatched embryos (Fig. 1A-C). Arrested unfertilized oocytes usually undergo meiotic maturation in the absence of sperm and become endomitotic, resulting in polyploidy and nuclear hypertrophy (see below) (*28*). We observed this phenotype across three different alleles of *rde-4*: *ne299* (A123→STOP), *uu53* (925 bp deletion), *pig51* (637 bp deletion) (Fig. 1A-D and fig. S1). We found that the fertility defects in *rde-4*(-) under heat stress stems from sperm failure, as crossing *rde-4*(-) to wild-type males rescued the phenotype (Fig. 1E).

**Fig. 1.**
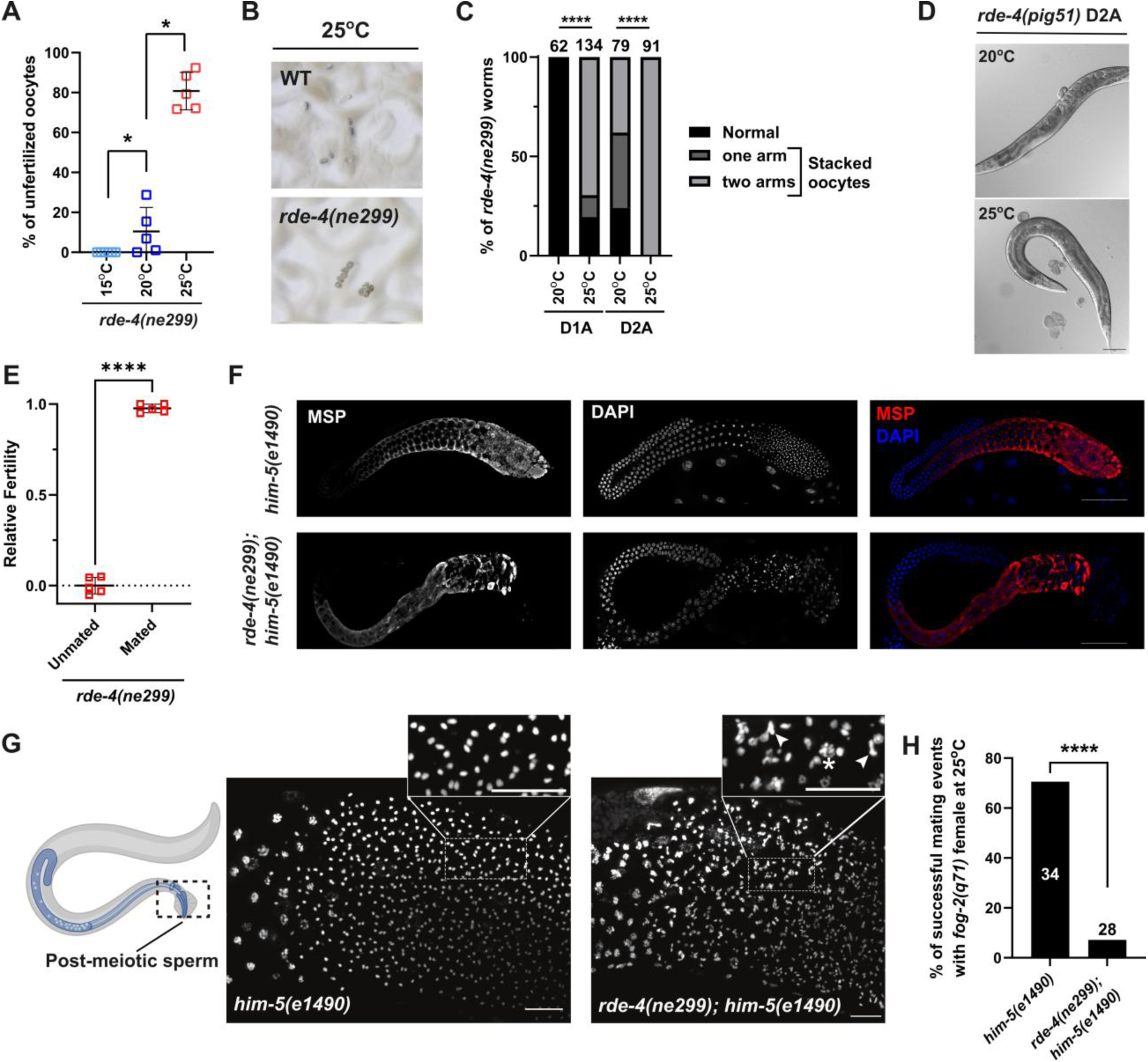
Loss of RDE-4 compromises sperm thermotolerance. (**A-D**) *rde-4(ne299)* shows temperature-sensitive fertility defects. (A) L4 larvae were transferred from 20 °C to 15 °C or 25 °C. The fertility of day 2 adults was scored. (B) At 25 °C, *rde-4(ne299)* animals lay unfertilized oocytes, which are morphologically distinct from fertilized embryos. (C) Unfertilized oocytes tend to accumulate and stack within the gonads of *rde-4(ne299)* at 25 °C. (D) *rde-4(pig51)* shows similar phenotypes to *rde-4(ne299)*. Scale bar = 100 μm. (**E**) The fertility defects of *rde-4(ne299)* at 25 °C is rescued by crossing with wild-type males. (**F**) MSP expression and localization is severely disrupted in male gonad in *rde-4(ne299); him-5(e1490)* at 25 °C. Scale bar = 50 μm. (**G**) Sperm from *rde-4(ne299); him-5(e1490)* exhibits chromosomal abnormalities. Arrows mark chromosomal bridges; asterisk indicates decondensation of DNA. Scale bar = 10 μm. (**H**) *rde-4(ne299); him-5(e1490)* males fail to mate with *fog-2(q71)* female at 25 °C. Error represents mean ± SD. * p ≤ 0.05; **** p < 0.0001.

Given the sperm defects observed in *rde-4*(-) hermaphrodites (Fig. 1A-E), we wondered whether *rde-4*(-) also plays a role in male sperm development. To enrich for males, we introduced a *him-5* mutation, which induces high frequency of X chromosome nondisjunction without compromising sperm morphology and functions (*29*). In *him-5(-)* males, major sperm protein (MSP), which is essential for sperm functions in nematodes, is packed into fibrous bodies and localized to membranous organelles; on the other hand, its organization is severely disrupted and MSP remain diffused in *rde-4(-); him-5(-)* males at 25 °C (Fig. 1F). In addition, we observed pronounced chromosomal abnormalities in spermatocytes from *rde-4(-); him-5(-)* males (Fig. 1G). Because *fog-2(-)* mutants lack self-sperm and rely on mating for reproduction, *fog-2(-)* females crossed with *him-5* males restored fertility. However, *rde-4(-); him-5(-)* males failed to sire progeny (Fig. 1H). Together, these results indicate that RDE-4, functioning in the endo-siRNA pathway, is required for the thermotolerance of sperm in both hermaphrodites and males.

## Neuron-to-germline communication affects sperm development

*C. elegans* contains a highly elaborate sRNA/AGO machinery, consisting of at least 19 AGOs that act sequentially to mediate potent gene silencing responses in diverse biological contexts (*30–32*). In the endogenous siRNA pathway, the ERI complex produces 26G sRNAs using mRNAs as templates. The 22G sRNAs are then loaded onto primary AGOs: ERGO-1 in oocytes and embryos, and ALG-3/4 in sperm, which recruit RNA-dependent RNA polymerases (RdRPs) RRF-1 and EGO-1, leading to the production of abundant secondary 22G sRNAs (*12*, *33*, *34*). In sperm, WAGO-10 and CSR-1a bind 22G sRNA and function downstream of ALG-3/4 to regulate spermatogenesis (*12*, *13*).

RDE-4 is required for the full production of 26G endo-siRNAs, although it is dispensable in certain cases (*25*, *35*, *36*). We found that siRNAs targeting spermatogenic genes are upregulated in *rde-4(-)* mutants grown at 25 °C compared to 20 °C (Fig. 2A). Notably, at 25 °C, upregulated siRNAs in *rde-4(-)* relative to wildtype are enriched for those targeting sperm genes, male-enriched genes, and known ALG-3/4-class sRNAs (Fig. 2B and fig. S2A).

**Fig. 2.**
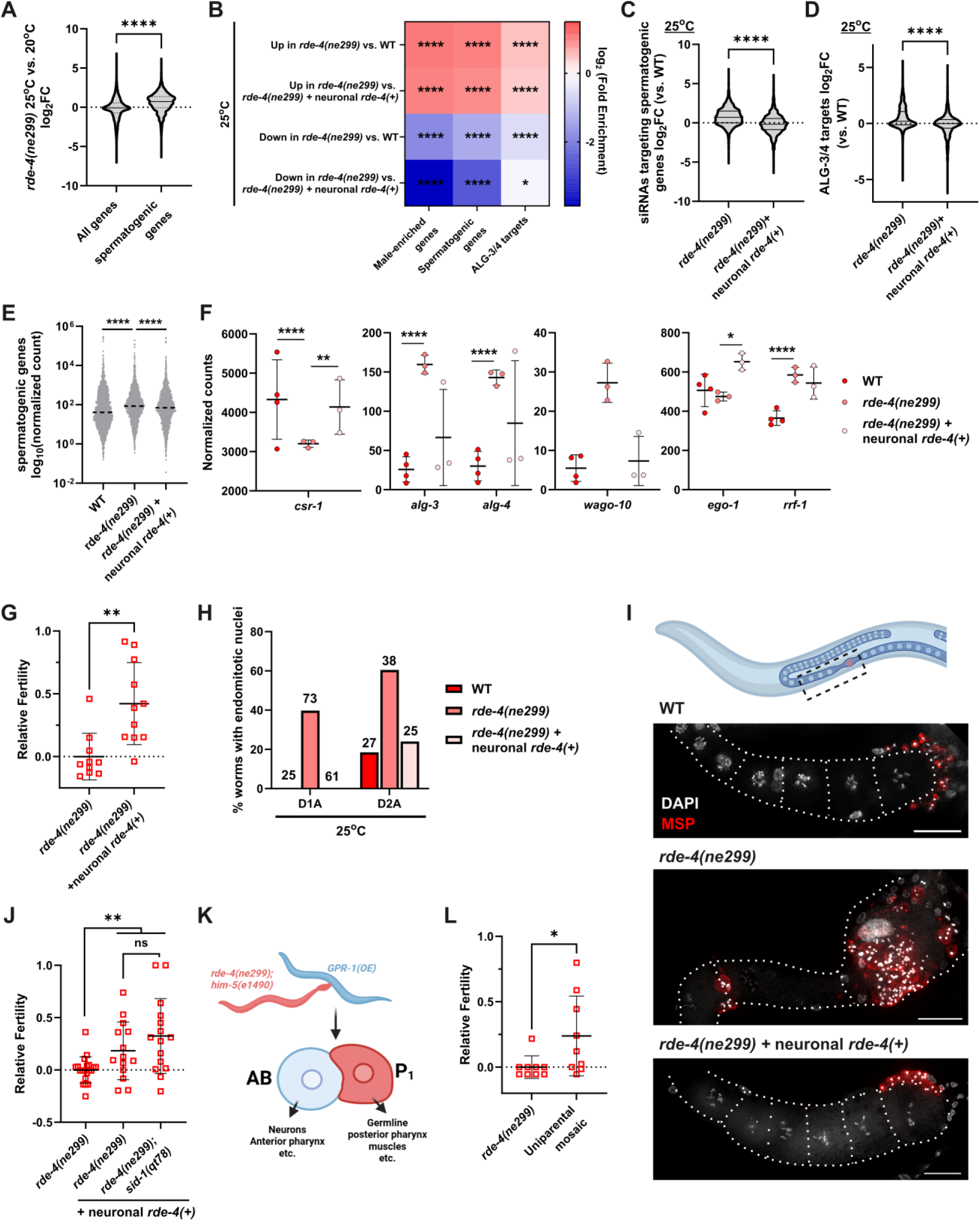
Neuronal RDE-4 promotes sperm thermotolerance. (**A**) siRNA targeting spermatogenic genes are upregulated in *rde-4(ne299)* grown at 25 °C compared 20 °C. (**B-D**) siRNAs targeting male-enriched genes, spermatogenic genes and ALG-3/4-class sRNAs are upregulated *in rde-4(ne299)* compared to wild type at 25 °C, which is rescued by neuronal *rde-4(+)* driven by *sng-1*/synaptogyrin promotor. (**E**) Spermatogenic transcripts are upregulated in *rde-4(ne299)* and restored by neuronal *rde-4(+)*. (**F**) Expression of AGO and RdRP genes are dysregulated in *rde-4(ne299)*, which is rescued by neuronal *rde-4(+)* in some cases. (**G-I**) Neuronal *rde-4(+)* partially rescues fertility defects in *rde-4(ne299)*. (G) Number of unfertilized oocytes laid were scored. (H) Percent of worms carrying endomitotic nuclei were scored. (I) *rde-4(ne299)* day 1 adult carries endomitotic nuclei and shows disrupted MSP expression at 25 °C. These phenotypes are rescued by neuronal *rde-4(+).* Scale bar = 20 μm. (**J**) The effects of neuronal *rde-4(+)* are not affected by the loss of *sid-1*. (**K**) Schematic of mosaic uniparental inheritance induced by GPR-1 overexpression. (**L**) Wild-type *rde-4* in the neurons of mosaic animals partially rescues fertility defects. Error represents mean ± SD. ns p > 0.05; * p ≤ 0.05; ** p < 0.01; **** p < 0.0001.

We recently demonstrated that neuronal RDE-4 may alter germline gene expression and trigger transgenerational behavioral changes (*37*). Intriguingly, in this study, we found that expressing *rde-4* in neurons as a single-copy MosSCI transgene under the control of pan-neuronal promotor (*pigSi3[Psng-1::rde-4:SL2:yfp]*) could largely restore the expression of sperm siRNAs, many of which are associated with ALG-3/4, in *rde-4(-)* mutants (Fig. 2B-D). ALG-3/4 has previously been shown to positively regulate its targets (*38*), and consistent with this, we observed aberrant upregulation of spermatogenic transcripts in *rde-4(-)* animals (Fig. 2E). In addition, we detected a widespread mis-regulation of several germline AGO, including *csr-1* and *wago-10*, and RdRP genes (Fig. 2F and fig. S2A). These expression patterns were at least partially rescued by neuronal *rde-4(+)* expression (Fig. 2B-F, and fig. S2B). Together, our findings indicate that neuronal RDE-4 regulates germline sRNA pathways cell-non-autonomously.

Strikingly, expression of neuronal *rde-4(+)* partially rescued the heat-induced fertility defects observed in *rde-4(-)* mutants (Fig. 2G-I). By performing smFISH and RNA-seq on isolated gonads, we previously confirmed that the neuronal *rde-4(+)* transgene is not mis-expressed in the germline (*37*). Importantly, similar to *rde-4(-)* mutants and in contrast to wild-type animals, *rde-4(-)* mutants carrying the neuronal *rde-4(+)* transgene are not responsive to exogenous dsRNA targeting *gfp* expressed in sperm (fig. S2C). Hence, these findings indicate that the observed effects are not artifacts caused by transgene mis-expression in the germline, but reflect bona fide cell-non-autonomous role of RDE-4 in regulating sperm heat tolerance. This neuron-to-sperm communication appears to be independent of SID-1, a conserved dsRNA-selective importer required for systemic RNAi (*39*) (Fig. 2J).

To substantiate our conclusion, we performed mosaic analysis by inducing uniparental isodisomy. To achieve this, we crossed *rde-4(ne299)* males to hermaphrodites overexpressing GPR-1, a conserved microtubule force regulator. This manipulation causes premature segregation of maternal and paternal chromosomes in the pre-cleavage embryos, resulting in non-Mendelian partitioning of genetic material between the first two blastomeres: AB and P_1_. In this system, the anterior AB contains exclusively maternally derived chromosomes, while the posterior P_1_ carries only paternally derived chromosomes (*40*, *41*). As nearly all neurons are derived from AB (*42*), they inherit the wild-type *rde-4* from the mother, whereas the germline, arise from the P_1_ lineage, carries the *rde-4(ne299)* mutation from the father (Fig. 2K). We found that these mosaic animals exhibit milder fertility defects compared to *rde-4(-)* mutants at 25 °C, supporting our earlier findings that neuronal sRNAs promote sperm thermotolerance (Fig. 2L). Importantly, this effect cannot be attributed to maternal provision, as *rde-4* homozygous mutants derived from heterozygous mothers still display severe fertility defects (fig. S2D).

## Oxygen sensing negatively impacts germline development

Previous studies showed that RDE-4 and endo-siRNAs are required for various neuronal processes, including olfactory adaption and dauer developmental decision (*43–46*). Importantly, we recently demonstrated that *rde-4(-)* mutants exhibits defective chemotaxis toward various volatile and soluble attractants at 25 °C, but not at 20 °C (*37*). This defect is partially rescued by neuronal expression of *rde-4(+)* (Data S1) (*37*). Hence, neuronal RDE-4 is important for the detection of various environmental cues, especially in the presence of heat stress.

Given the roles of RDE-4 in sensory signaling, we tested whether blocking sensory perception affects sperm development. Interestingly, eliminating *cmk-1*, a gene encoding the homolog of CaMKI which modulates sensory gene expression, partially rescues *rde-4(-)* sterility at 25 °C (Fig. 3A). We observed similar effects when we knockout *tax-2*, which encodes a subunit of cyclic nucleotide (cGMP)-gated channel that is required for many sensory responses (Fig. 3B, C). These results indicate that the sensory inputs normally inhibit sRNA-mediated sperm heat tolerance.

**Fig. 3.**
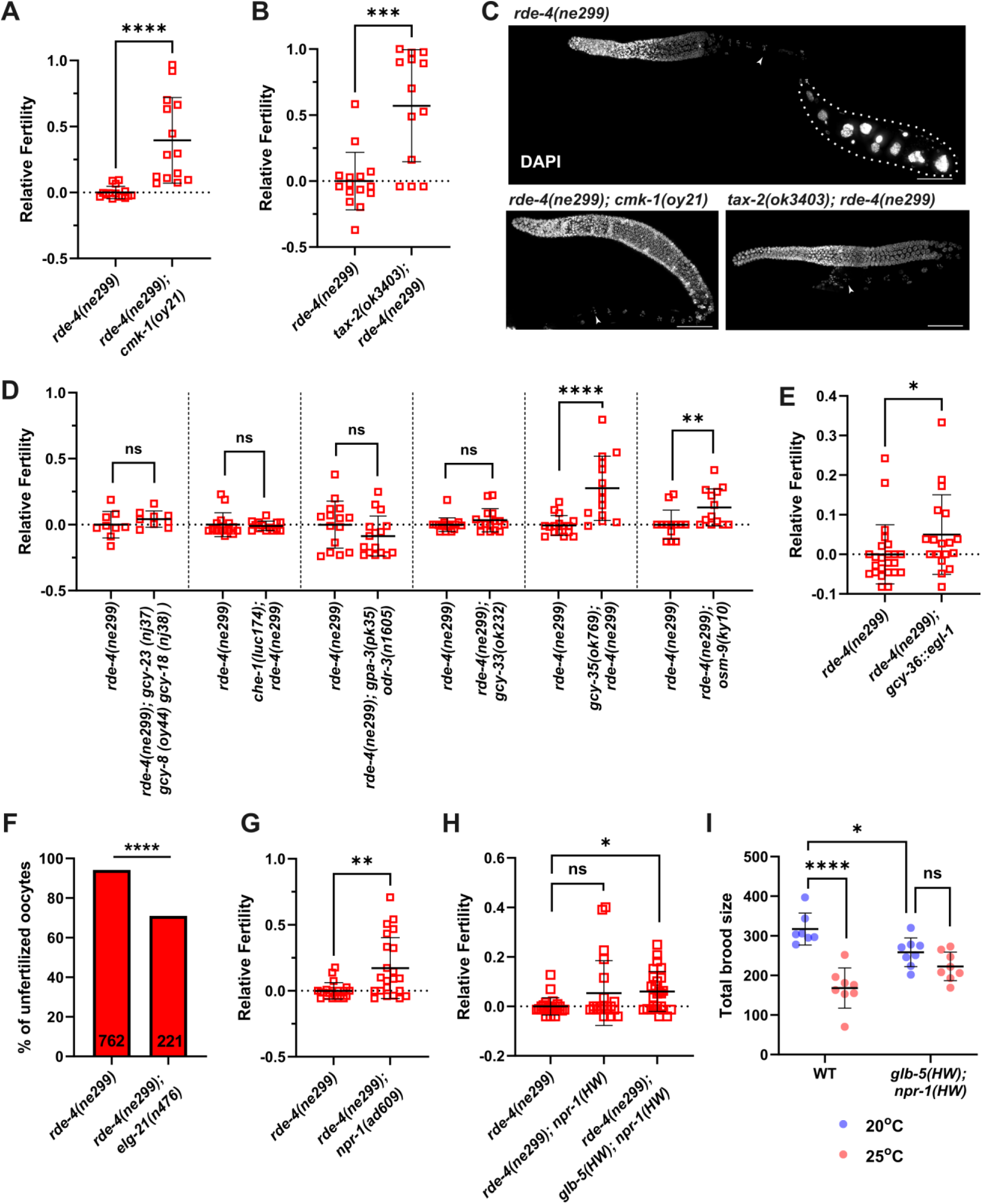
Oxygen-sensing neurons inhibit reproduction. (**A-C**) Broad inhibition of sensory perception by knocking out *cmk-1* or *tax-2* rescues the fertility defects of *rde-4(ne299).* (C) Endomitotic oocytes in *rde-4(ne299)* are highlighted. Arrows indicate the -1 oocytes, located closest to the spermatheca. Scale bar = 50 μm. (**D**) Inhibiting oxygen perception, but not the other sensory modalities, improves *rde-4(ne299)* fertility. Knocking out *gcy-35* or *osm-9* rescues *rde-4(ne299)* phenotype. (**E**) Genetically ablating URX, AQR, and PQR neurons partially rescues *rde-4(ne299)* sterility. (**F**) Blocking neuropeptide processing rescues *rde-4(ne299)* sterility. Note that *egl-2(n476)* has egg-laying defects independent of fertility. (**G**) EMS-induced *npr-1(ad609)* loss-of-function mutation rescues *rde-4(ne299)* sterility. (**H-I**) Ancestral (HW) alleles of *npr-1* and *glb-5* partially restore *rde-4(ne299)* fertility. *glb-5(HW); npr-1(HW)* double mutants do not show heat-induced loss of fertility. Error represents mean ± SD. ns p > 0.05; * p ≤ 0.05; ** p < 0.01; p *** < 0.001; **** p < 0.0001.

Next, we asked which sensory modalities might regulate sperm development. Given that sterility in *rde-4(-)* mutants is temperature-sensitive, we first eliminated three guanylyl cyclases (GCs) – GCY-8, CGY-18, and GCY-23 – which function specifically in AFD neurons, the primary thermosensory neurons in *C. elegans* (*47*). While we previously showed that removing *gcy-8/18/23* downregulates AGO genes and disrupts heritable gene silencing in the germline (*48*), we found that blocking heat sensation had no effect on the fertility of *rde-4(-)* mutants. Similarly, impairing gustatory signaling by removing CHE-1, a C2H2-type zinc finger TF required for specification of the ASE gustatory neurons (*49*, *50*), or disrupting general chemosensation by knocking out the G protein α subunits GPA-3 and ODR-3 (*51*), also did not impact fertility (Fig. 3D). In addition, eliminating MEC-8, which plays a role in amphid cilia fasciculation and mechanosensory neuron development (*52*, *53*), had no observable effect (fig. S3A).

*C. elegans* has strong behavioral responses to the gases oxygen and carbon dioxide, which are highly variable in its natural habitats such as soil, compost heaps, and rotting substrates. Environmental oxygen is primarily detected by the AQR, URX, and BAG neurons in the head, and the PQR neuron in the tail, with additional contribution from other neurons (*54–56*). Mitochondrial activity modulates cellular oxygen consumption, affecting behavioral and physiological response to environmental oxygen levels (*57*, *58*). We found that genes downregulated in *rde-4(-)* mutants at 25 °C compared to those at 20 °C are significantly enriched for mitochondrial processes (fig. S3B). We observed a similar pattern of mitochondrial gene downregulation when comparing *rde-4(-)* mutants to wild-type animals at 25 °C (fig. S3C and Data S1).

We found that deletion of *gcy-33*, an oxygen receptor gene expressed in BAG neurons (*56*, *59*), did not affect the fertility of *rde-4(-)* mutants (Fig. 3D). In contrast, loss of *gcy-35*, which encodes a heme-containing oxygen sensor expressed in the URX, AQR, PQR, SDQ, ALN, and PLN neurons that promote hyperoxia avoidance, rescues sterility of *rde-4(-)* mutants (Fig. 3D and fig. S3D). Genetically ablating URX, AQR, and PQR by expressing the cell-death activator gene *egl-1* driven by the *gcy-36* promoter causes a modest but significant rescue of *rde-4(-)* sterility (Fig. 3E).

TRPV sensory channel OSM-9 acts in the polymodal nociceptive ASH neurons to regulate avoidance of high osmolarity, chemical repellents and touch, and also acts in non-nociceptive cell types to mediate olfactory responses and sensory adaptation (*60–62*). Of note, OSM-9 has been shown to function in ASH and serotonergic ADF neurons to promote hyperoxia avoidance (*54*). We found that knocking out *osm-9* modestly rescues *rde-4(-)* phenotype (Fig. 3D). However, removing the tryptophan hydroxylase enzyme required for serotonin biosynthesis TPH-1 does not affect *rde-4(-)* fertility (fig. S3E), nor does genetic ablation of ASH neurons alone (fig. S3F). Together, these results indicate that multiple oxygen-sensing neurons function synergistically to regulate sRNA-mediated germline development. This neuron-to-germline communication is likely mediated by neuropeptides, as removing carboxypeptidase E orthologue EGL-21, which is required to process endogenous neuropeptides, partially restores *rde-4(-)* fertility (Fig. 3F).

Hypoxia exposure, for example living at high altitude, has long been known to negatively impact male fertility in animals (*63*). Here, we show that mere perception of oxygen – without changing oxygen levels – can antagonize sperm development and fertility modulated by small RNA pathways. We propose that inhibiting oxygen-sensing neurons may mimic cues associated with active bacterial growth that create oxygen sinks, suggestive of conditions favorable for reproduction in *C. elegans*.

Neuronal signaling pathways, especially the sensory system, exhibit extensive evolutionary plasticity and may be subjected to positive selection in response to changing habitats (*64*, *65*). In the canonical *C. elegans* “wild-type” strain (N2), laboratory domestication led to the fixation of a gain-of-function allele in the neuropeptide Y homolog *npr-1* and a duplication/insertion in globin gene *glb-5*, both of which alter the animals’ response to oxygen and carbon dioxide. These laboratory-derived alleles are absent in LSJ1, a sister strain derived from N2 at least six years before its cryopreservation in 1969 (*66*).

In N2, the gain-of-function NPR-1(215V) downregulates the activities of oxygen-sensing AQR, PQR, and URX neurons, thereby suppressing hyperoxia avoidance. NPR-1(215V) animals do not respond to drops in oxygen levels and are solitary feeders, dispersing across bacterial lawn. In contrast, wild isolates such as CB4856 from Hawaii (abbreviated as HW), which carry NPR-1(215F), or animals with an *npr-1* loss-of-function allele, slow their movement in response to reduced oxygen and exhibit social feeding, aggregating along the bacterial lawn border where oxygen levels are lower (effective oxygen concentration of ∼12.8% vs. ∼17.1% in the center of the lawn) (*55*).

Interestingly, we found that a null mutation of *npr-1* (an EMS-induced allele) rescues the fertility defects of *rde-4(-)* mutants, likely due to social aggregation in low oxygen environments (Fig. 3G). Although *npr-1(HW)* alone does not markedly affect *rde-4(-)* fertility, *glb-5(HW); npr-1(HW)* double mutant significantly rescues *rde-4(-)* phenotype (Fig. 3H). Consistent with previous findings that these two laboratory-derived alleles confer significant fitness advantages in standard laboratory environment, *glb-5(HW); npr-1(HW)* mutants show a lower brood size compared to wildtype at 20 °C (Fig. 3I) (*67*, *68*). At 25 °C, the brood size of wild-type animals drops significantly owning to sperm dysfunction (*69*); we found that this decline is not evident in *glb-5(HW); npr-1(HW)* (Fig. 3I), suggesting that the ancestral alleles of *npr-1* and *glb-5* promote reproductive resilience under heat stress – a major challenge in their natural habitat.

Hence, our results indicate that laboratory domestication alters oxygen-sensing circuit and foraging behaviors, which subsequently impact sRNA-mediated germline development under heat stress. However, we note that knocking out *rde-4* in LSJ1 and CB4856 causes severe loss of fertility, as observed in N2 (Fig. S3G), suggesting other natural genetic variants in these strains may act to affect germline development and/or sRNA pathways (*70–73*). Indeed, AGO genes and sRNA pathways are evolving rapidly, exhibiting extensive intraspecies variation that may have broad impacts on germline gene regulation (*71*, *74*). Additionally, other laboratory-derived alleles have been shown to affect sperm development (*72*, *75*), including a deletion in *nurf-1*, which encodes the orthologue of the BPTF subunit of the NURF chromatin remodeling complex, that arose in the LSJ1 lineage (*75*).

## Neuronal signaling influences genome stability in the germline

We note that, although mating largely rescues the fertility defect in *rde-4(-)* mutants, it does not fully restore brood size to wild-type levels, suggesting additional, non-sperm-origin defects (Fig. 4A). To investigate further, we examined RAD-51/RecA, which localizes to the double-strand break (DSB) repair loci (*76*, *77*). In wild-type gonads, RAD-51 appears as distinct foci mostly in the leptotene/zygotene and pachytene stages coincide with the onset of meiosis (Fig. 4B, C). In contrast, at 25 °C, but not at 20 °C, *rde-4(-)* mutants exhibited an elevated frequency of DSBs throughout the germline, including the mitotic proliferative stem cells, indicating compromised genome integrity under heat stress (Fig. 4D and fig. S4A). Remarkably, consistent with our earlier findings, this defect was rescued by single-copy neuronal expression of *rde-4(+)* (Fig. 4E-G). Moreover, eliminating *cmk-1* or *tax-2* reduces the number of RAD-51 foci in *rde-4(-)* mutants (Fig. 4H and fig. S4B). Finally, loss of *gcy-35* rescued the DSBs in *rde-4(-)* (Fig. 4I-K), further supporting our conclusion that neuronal activity – particularly the oxygen-sensing circuit – plays an important role in regulating reproductive robustness. Perception of oxygen levels may promote germline maintenance as low oxygen levels correlate with bacteria-rich environments and reproductive permissive niche (*55*, *78*).

**Fig. 4.**
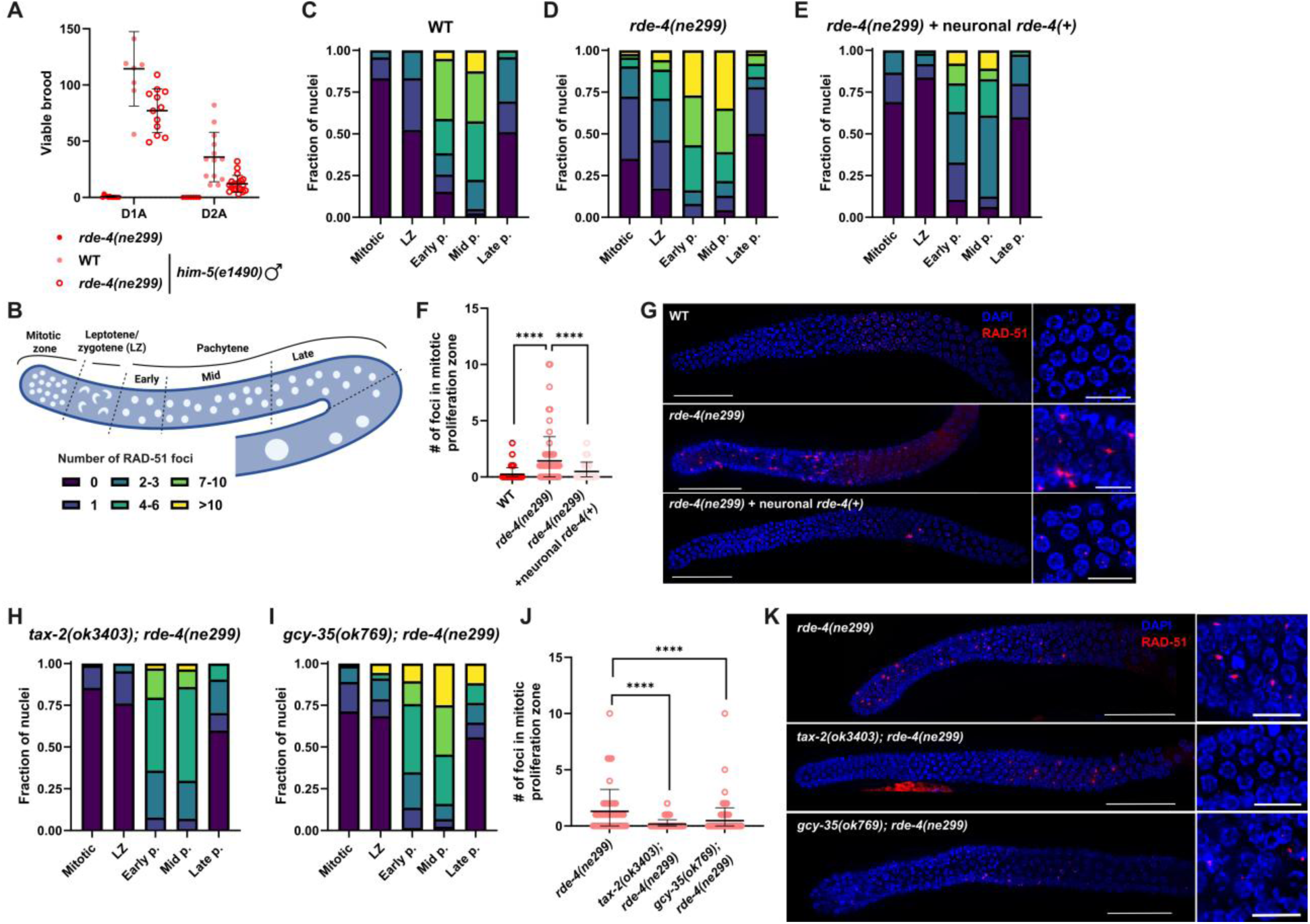
Neuronal RDE-4 and impaired oxygen-sensing neurons promote germline genome stability. (**A**) Crossing *rde-4(ne299)* hermaphrodites to *him-5(e1490)* males does not completely restore brood size at 25 °C. (**B-G**) Neuronal *rde-4(+)* rescues DSBs in *rde-4(ne299)* at 25 °C. (B) Diagram showing the five regions where RAD-51 foci were quantified. (C-E) Stacked bar charts showing the fraction of nuclei in each region, color-coded by the number of RAD-51 foci, in wildtype, *rde-4(ne299)*, and *rde-4(ne299) + neuronal rde-4(+)* at 25 °C. (F) Wild-type and *rde-4(ne299) + neuronal rde-4(+)* animals contain fewer number of DSBs in the mitotic germ cells compared to *rde-4(ne299)*. (G) Whole gonads stained with RAD-51 (red) and DAPI (blue). Scale bar = 50 μm. Right panels show magnification of mitotic nuclei. Scale bar = 10 μm. (**H-K**) Loss of oxygen perception rescues DSBs in *rde-4(ne299).* (H, I) Stacked bar charts showing the number of RAD-51 foci in the germ cells across different regions in *tax-2(ok3403); rde-4(ne299)* and *gcy-35(ok769); rde-4(ne299).* (J) *tax-2(ok3403); rde-4(ne299)* and *gcy-35(ok769); rde-4(ne299)* contain fewer number of DSBs in the mitotic germ cells compared to *rde-4(ne299)* at 25 °C. (K) Whole gonads stained with RAD-51 and DAPI in *rde-4(ne299)*, *tax-2(ok3403); rde-4(ne299)* and *gcy-35(ok769); rde-4(ne299).* Scale bar = 50 μm. Right panels show magnification of mitotic nuclei. Scale bar = 10 μm. Error represents mean ± SD. **** p < 0.0001.

Regulatory sRNAs can exert cell non-autonomous effects via microvesicles or exosomes, influencing diverse biological processes in both health and disease (*79*, *80*). The intercellular transfer of RNAs and other molecular cargoes mediates soma–germline interactions from plants to mammals, regulating germline gene expression (*81–83*). In this study, we report that neuronal sRNAs can regulate reproduction through a neuroendocrine-like signaling mechanism that depends on neuropeptides, rather than on direct RNA transfer. This neuron-to-germline communication may, in some cases, initiates a lasting transgenerational epigenetic memory (*48*). Hence, we propose that sRNAs integrate sensory information, promote developmental plasticity and reproductive adaptation, and potentially facilitate epigenetic buffering in fluctuating environments (*84–87*). The sensory regulation of sRNA-mediated germline mortality may exert a profound influence on the evolutionary trajectories of animals.

## Materials and Methods

### Worm cultivation

*C. elegans* strains were maintained on standard Nematode Growth Medium (NGM) plates seeded with *E. coli* OP50 at 20 °C, unless stated otherwise. A complete list of strains used in this study is provided in Table S1.

### Fertility assay

L4 animals, identified by the characteristic white crescent surrounding the developing vulva, were transferred to 25 °C. After ∼24 hours, individual day 1 adults were moved to fresh NGM plates seeded with a small (∼30 µL) drop of OP50 that had been allowed to dry for 1-2 days. About 12 hours later, the number of laid eggs and unfertilized oocytes was quantified. Samples with fewer than 20 total eggs and unfertilized oocytes were excluded from analysis. Relative fertility was calculated as: 1 – (% unfertilized oocytes / mean % unfertilized oocytes in control *rde-4(-)* animals from the same experiment). In all cases, the investigators were blinded to the genotypes.

### RNA isolation

Total RNA was isolated from day 1 adults (∼72 hours at 20°C following L1 synchronization) using standard phenol-chloroform method with TRIzol^TM^ (Invitrogen). RNA quality was accessed using Agilent 4150 BioAnalyzer instrument and High Sensitivity RNA ScreenTapes.

### RNA-seq

To sequence the mRNA, RNA libraries were prepared using NEBNext® Ultra II Directional RNA Library Prep Kit for Illumina® coupled with NEBNext® Poly(A) mRNA Magnetic Isolation Module. The cDNA libraries were pooled and paired-end sequencing was performed on the Nextseq 2000 platform.

For sRNA sequencing, we treated the RNA samples with RNA 5′ Polyphosphatase (epicentre) and the libraries were prepared using NEBNext® Multiplex Small RNA Library Prep Set for Illumina (New England Biolabs) or TruSeq Small RNA Library Prep Kit (Illumina) according to the manufacturer’s protocol. RNA ranging from ∼140 to 160 nt was size-selected by gel extraction using 4% agarose E-Gel (Life Technologies). The pooled samples were the sequenced using the Illumina HiSeq 2500 instrument.

### Bioinformatics

All bioinformatic analyses were performed using RNAlysis (*88*). For sRNA-seq data, sRNA reads were aligned to PRJNA13758 ce11 genome assembly using ShortStack (*89*) and aligned anti-sense reads were quantified using FeatureCounts (Data S2) (*90*). For mRNA, we pseudo-aligned reads using Kallisto (Data S3) (*91*). We then performed differential expression analysis using DESeq2 (*92*). For gene set enrichment analysis, log_2_ (fold enrichment) scores were computed and the FDR for enrichment was calculated using 10,000 random gene sets identical in size to the tested group. For Gene Ontology (GO) and KEGG pathways analyses, Fisher’s Exact tests were performed.

### CRISPR/Cas9 gene editing

CRISPR/Cas9 was performed as previously described (*93*). To generate *rde-4* deletion, we used two crRNAs: ACTTGGGGCACTGTCGAACT and ATCTCTGGAATCATATGATA. The following homology-directed repair (HDR) donor was used: AGGCTGCTAAGGCTGTCTATCAAAAGACGCCAACTATATGGGTATGCCTCCAAATAATTGTAGTTAATAT. This generates 637 bp deletion in *rde-4*. The results strains were backcrossed at least two times to remove potential background mutations.

### Antibody and DAPI staining

Dissected gonads were fixed with 3.5% formaldehyde for 10 mins and washed at least twice with PBS-T (0.1% Tween-20 in PBS buffer). Tissues were then permeabilized in 0.25% PBS-Triton for 20 mins, followed by blocking in 0.5% BSA in PBS-T for 1 hour with gentle shaking. Next, primary antibodies – mouse anti-MSP (Developmental Studies Hybridoma Bank) and rabbit anti-RAD-51 (gift from Nicolas Silva) – were diluted 1:1000 in PBS-T and applied for 1 hour with gentle agitation. After two washes with PBS-T, secondary antibodies (Jackson ImmunoResearch anti-rabbit and anti-mouse; 1:200 in PBS-T) were added and incubated for 1 hour. Samples were then washed for two more times and were incubated with DAPI (0.02% in PBS-T) for 10 mins, followed by two more washes with PBS-T. Finally, samples were mounted on glass slides in VECTASHIELD® Antifade Mounting Medium (Vector Laboratories, Cat# H-1000) prior to imaging.

### Statistical analysis

Statistical analyses were conducted using R software v4.2.3 and GraphPad Prism 10. For comparisons between two groups, non-parametric unpaired Wilcoxon rank sum tests were used. For experiments involving multiple groups, Kruskal-Wallis test followed by pairwise Wilcoxon rank-sum tests were performed. Multiple comparison corrections using the Benjamini–Hochberg method were applied where appropriate.

## Acknowledgments

We thank Rutwik Bardapurkar and Itai Reiger for their assistance with experiments. We thank Cori Bargmann (Rockefeller University) for providing introgressed strains carrying HW alleles of *npr-1* and *glb-5*. Some graphics were created with Biorender.com. We are grateful to WormBase for providing valuable data and resources. Some strains were provided by the CGC, which is funded by NIH Office of Research Infrastructure Programs (P40 OD010440).

## Author contributions

Conceptualization: CKE, OR

Methodology: CKE, HA

Investigation: CKE, HA, SW, AM, GT, SA, HG, RP, OA

Visualization: CKE, HA

Funding acquisition: YBT, OR

Supervision: YBT, OR

Writing – original draft: CKE

Writing – review & editing: CKE, HA, YBT, OR

## Data and materials availability

All data are available in the main text or the supplementary materials. All NGS data are available through GEO, accession number GEO: XX.

## Supplementary Materials

**Fig. S1.**
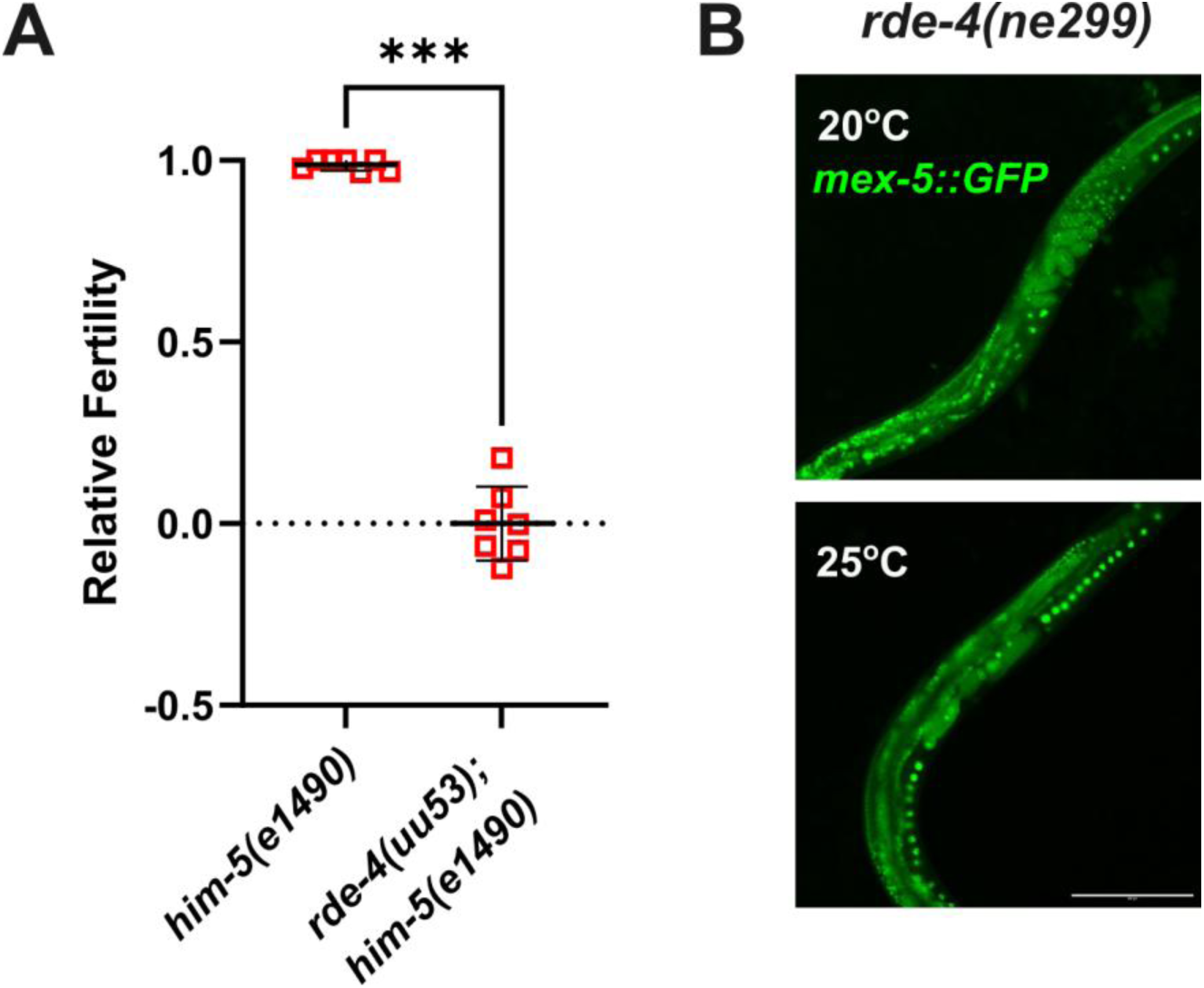
RDE-4 promotes reproduction at high temperature. (**A**) *rde-4(uu53)* mutants exhibit severe loss of fertility at 25 °C. (**B**) *rde-4(ne299)* day 2 adult shows accumulation of unfertilized oocyte stacked in the gonad at 25 °C, but not at 20 °C. The germline is marked by *gfp* driven by *mex-5* (RNA-Pol II) promotor. *** p < 0.001.

**Fig. S2.**
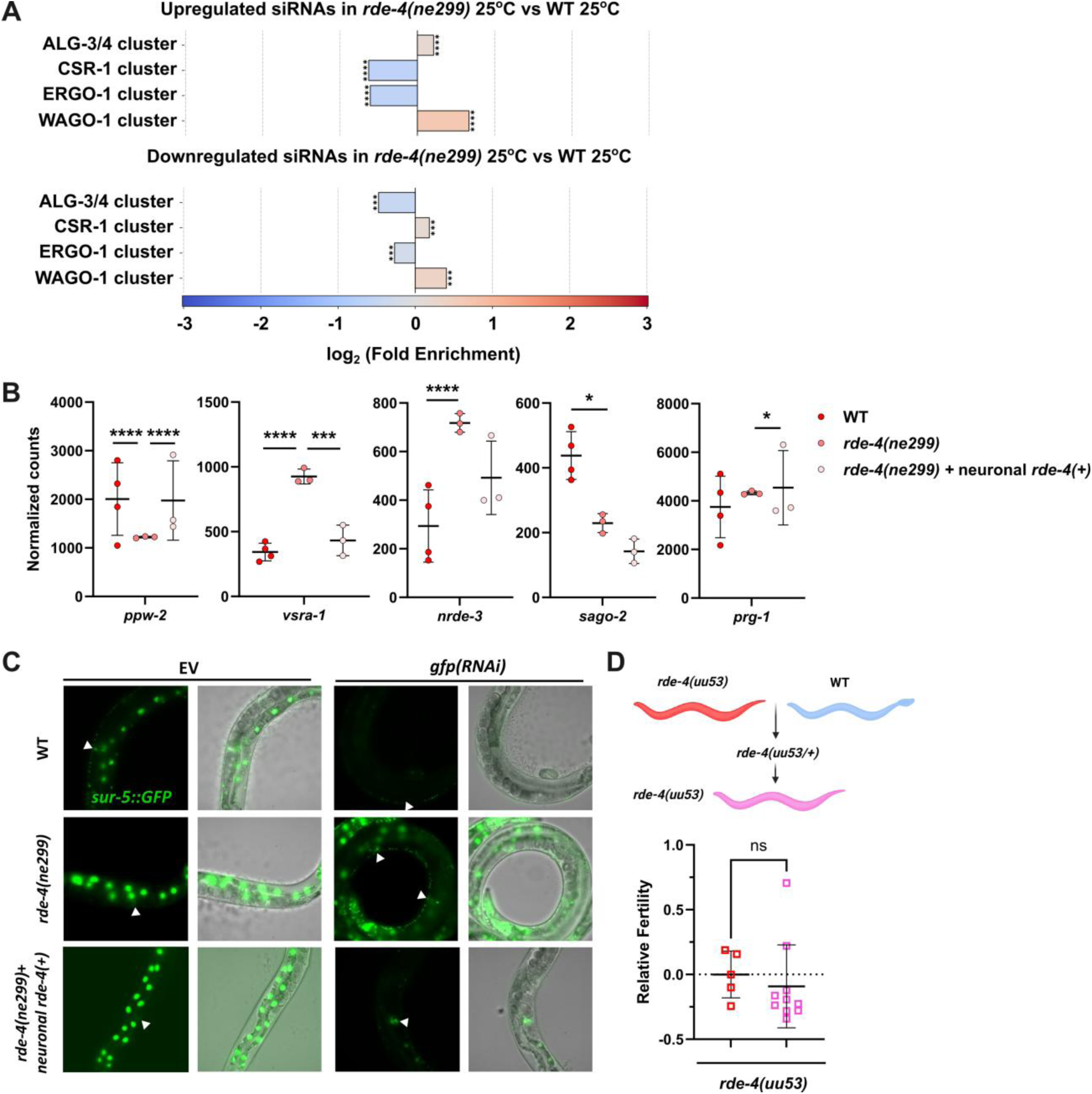
Neuronal RDE-4 promotes sperm development. (**A**) Different classes of endo-siRNAs are differentially expressed in *rde-4(ne299)*. ALG-3/4-class siRNAs tend to be upregulated, whereas CSR-1–class siRNAs tend to be downregulated in *rde-4(ne299)* compared to wild type. (**B**) Misexpression of AGO genes in *rde-4(ne299)* is rescued by neuronal *rde-4(+)* in some cases. (**C**) Neuronal *rde-4(+)* does not rescue RNAi-defective phenotype of *rde-4(ne299)*. Arrows indicate spermatheca. (**D**) Homozygous *rde-4(uu53)* segregated from heterozygous mothers show severe fertility defects. ns p > 0.05; * p ≤ 0.05; *** p < 0.001; **** p < 0.0001.

**Fig. S3.**
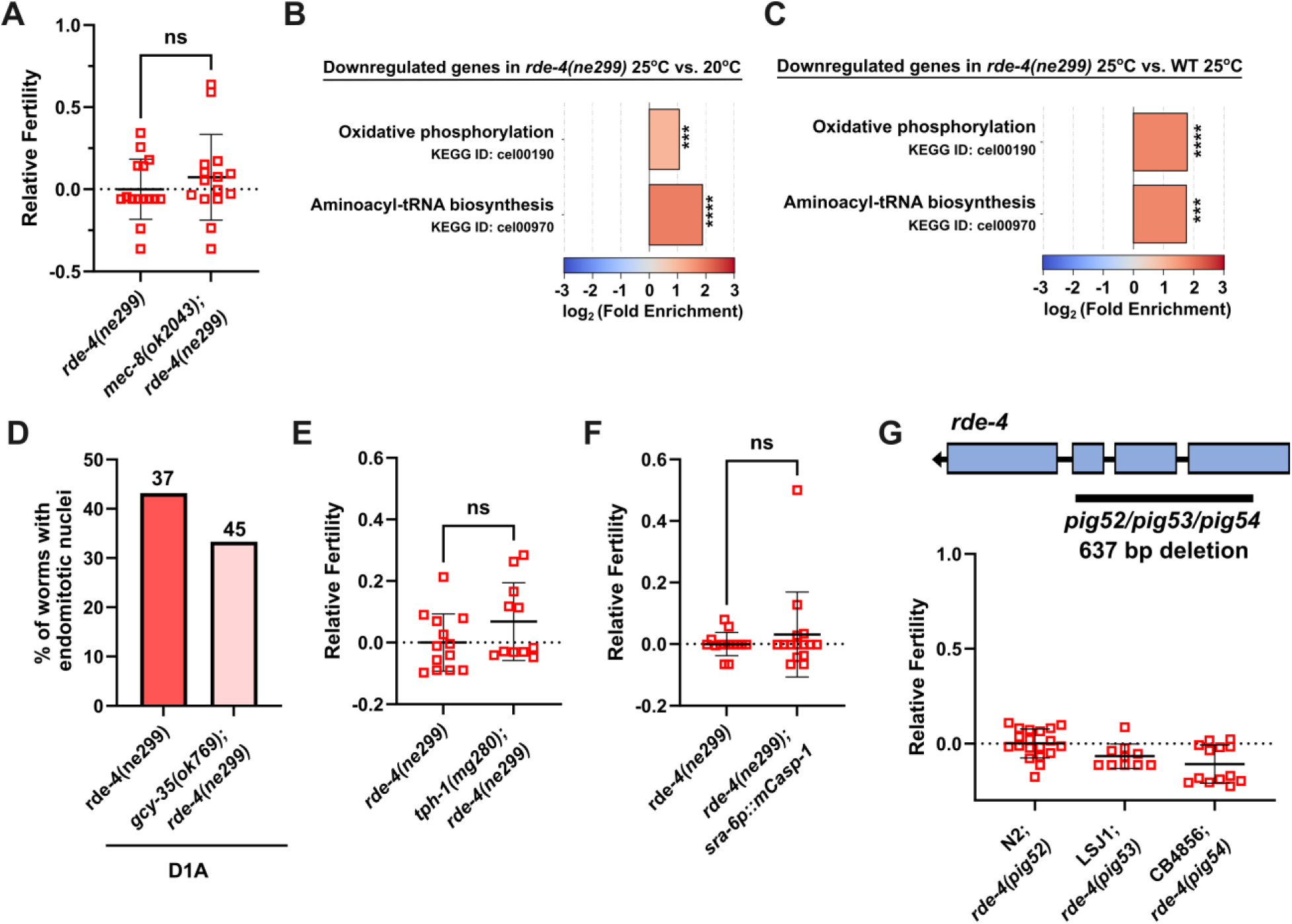
Neuronal sensory affects fertility. (**A**) Knocking out *mec-8* does not affect *rde-4(ne299)* fertility. (**B-C**) KEGG pathway analysis reveals that mitochondrial genes tend to be downregulated in *rde-4(ne299)* grown at 25 °C compared to either *rde-4(ne299)* at 20 °C or wild type at 25 °C. (**D**) Knocking out *gcy-35* reduces the accumulation of endomitotic nuclei. (**E**) Eliminating *tph-1* does not rescue *rde-4(ne299)* sterility. (**F**) Genetically ablating ASH neurons does not impact *rde-4(ne299)* fertility. (**G**) Deleting *rde-4* in CB4856 and LSJ1 causes severe loss of fertility at 25 °C, similar to that observed in N2. ns p > 0.05; **** p < 0.0001.

**Fig. S4.**
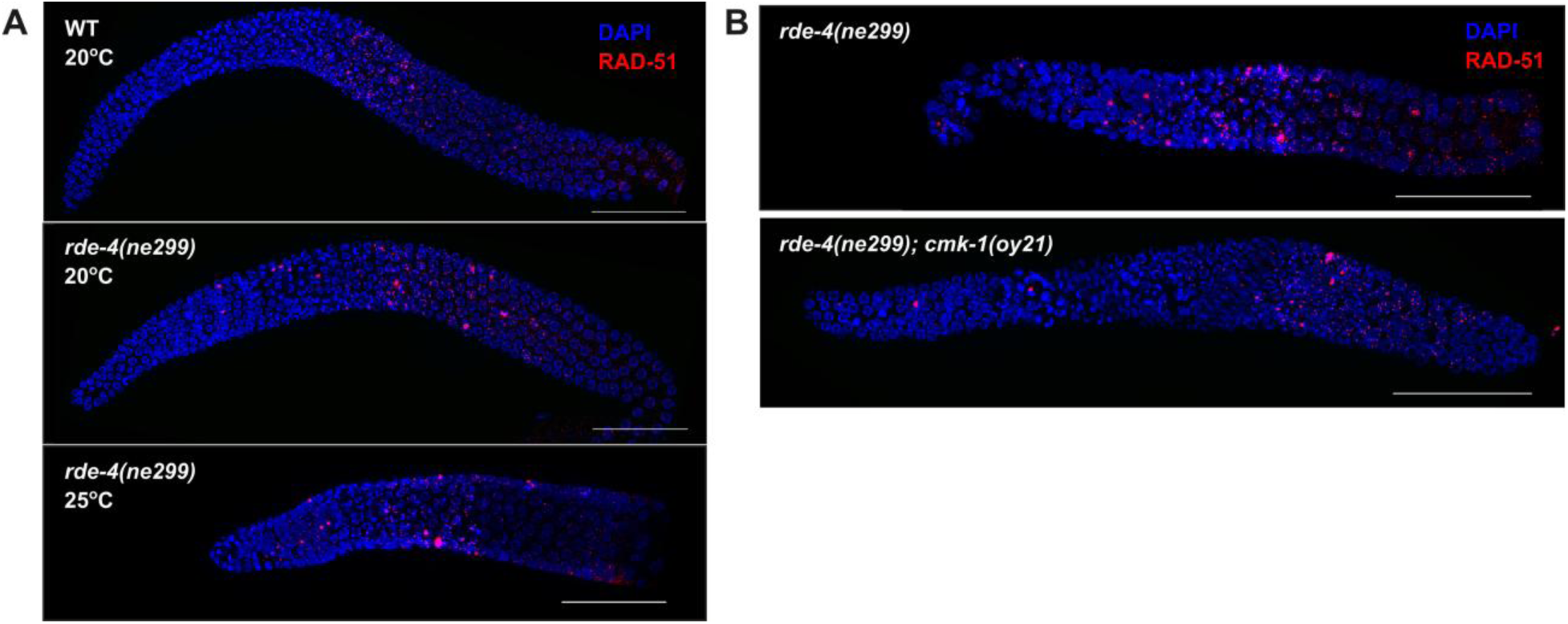
Neuronal sensory affects germline integrity. (**A**) *rde-4(ne299)* grown at 25 °C, but not 20 °C, shows increased DSBs in the gonad. (**B**) Knocking out *cmk-1* reduces DSBs in *rde-4(ne299)* at 25 °C. Scale bar = 50 μm.

**Table S1.**
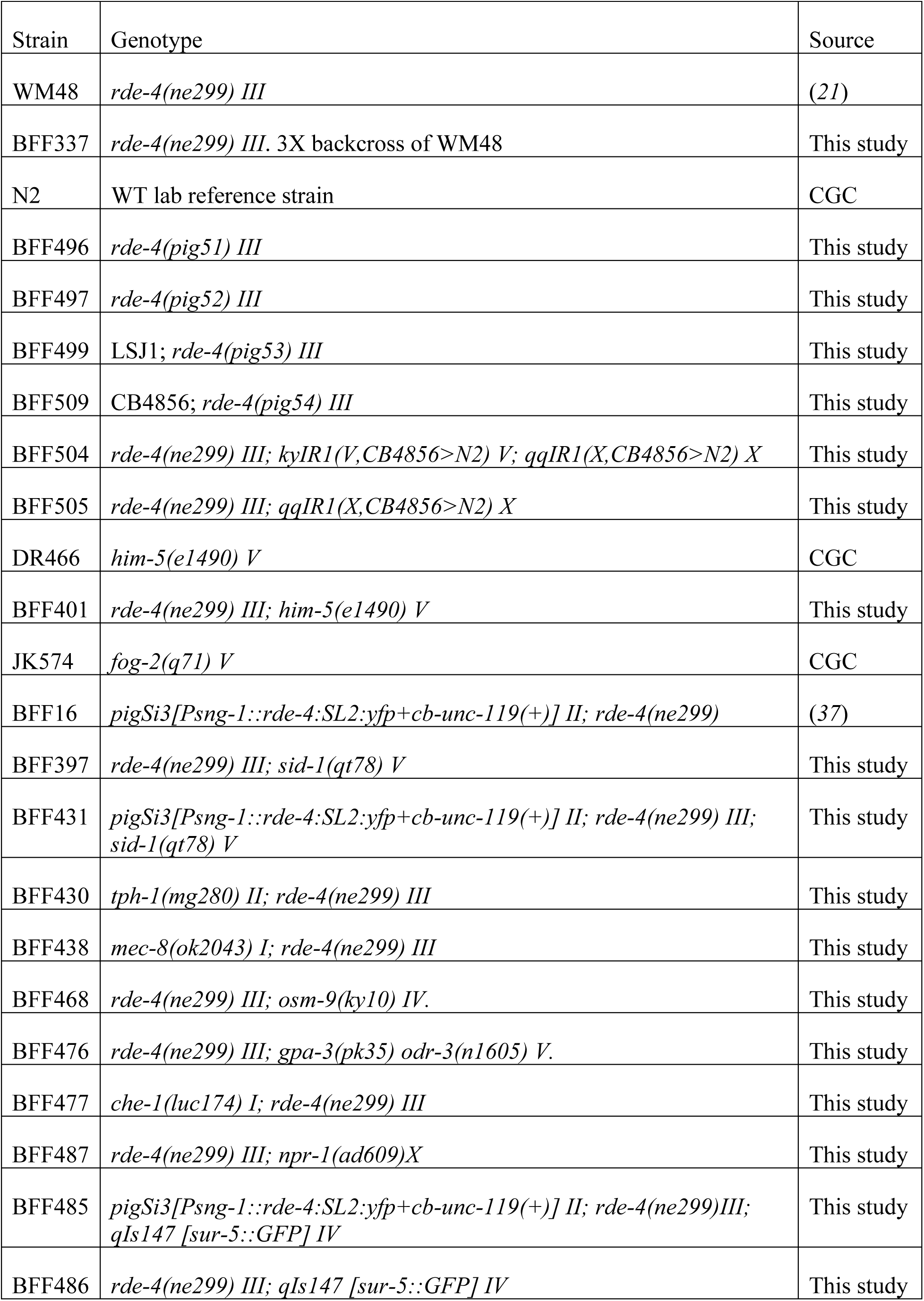
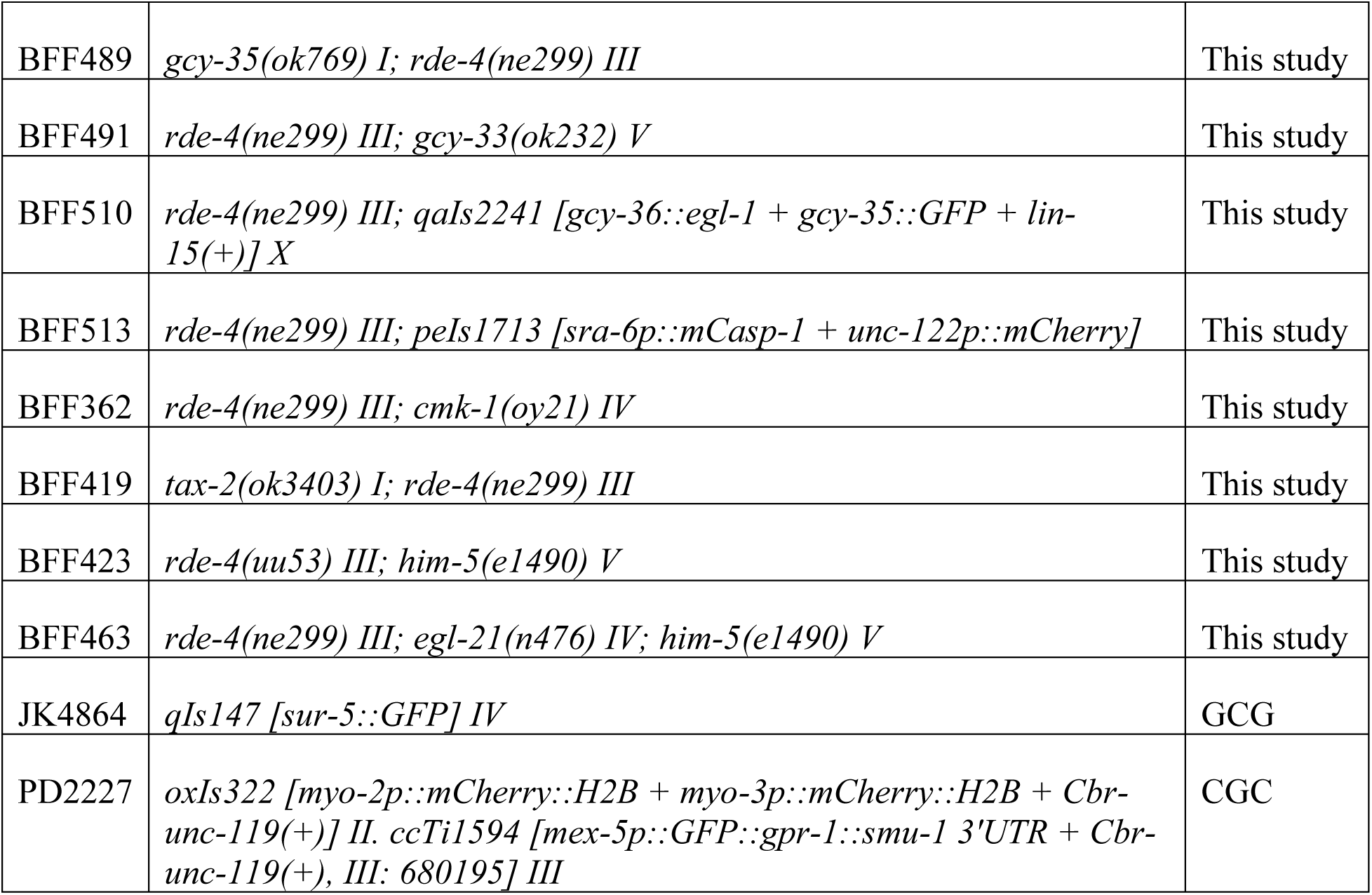
Worm strains used in this study.

